# Isoflavones impair response to anti-PD1 therapy in murine breast cancer models, irrespective of dietary fiber and fecal short chain fatty acid levels

**DOI:** 10.64898/2025.12.05.692375

**Authors:** Fabia de Oliveira Andrade, Kerrie B. Bouker, Melike Ozgul-Onal, Lu Jin, Idalia Cruz, William Helferich, Audrey Gao, Karla Andrade, Vivek Verma, Christopher Staley, Patricia L Foley, Leena Hilakivi-Clarke

**Affiliations:** The Hormel Institute, University of Minnesota, Austin, MN 55912, USA; Georgetown University Medical Center, Washington DC, 20057, USA; Department of Histology-Embryology, Medicine, Mugla Sitki Kocman University, Mentese-Mugla, Turkey; Department of Food Science and Human Nutrition, University of Illinois Urbana-Champaign, Urbana, IL 61801, USA; Department of Biochemistry and Pharmacology, Federal University of Piaui, Teresina, PI 64. 049-550, Brazil; Department of Surgery, Medical School, University of Minnesota, Twin Cities Campus, MN 55455, USA; Depatment of Food Science and Nutrition, University of Minnesota, St. Paul, MN 55108

**Keywords:** Microbiota assessable carbohydrates (MACs), genistein, anti-PD1, tamoxifen, gut microbiome, short chain fatty acids (SCFAs), tumor immune genes

## Abstract

**Background:** Fermentable dietary fibers, also called microbiota-accessible carbohydrates (MAC), and the consequent increase in fecal short-chain fatty acids (SCFAs) are linked to improved responsiveness to immune checkpoint blockade (ICB) therapy in human and mouse studies. However, experimental diets high in MAC also often contain estrogenic isoflavones, which may counter fiber’s beneficial effects by causing immunosuppression.

**Methods:** We studied the effects of feeding female C57BL/6Tac mice low-MAC (AIN93G), low-MAC supplemented with isoflavone genistein, high-MAC (5V5M) or high-MAC isoflavone (high-MACi; 5058D) diet on their gut microbiome and response to anti-PD1 therapy against E0771 allografted triple negative breast cancer (TNBC) and 7,12-dimethylbenz[a]anthracene (DMBA)-initiated estrogen receptor α positive (ERα+) mammary tumors. We also determined whether blocking ERα with tamoxifen (TAM) impacted responsiveness to anti-PD1 therapy in mice fed different diets. The effect of diet and treatments on immune cell signaling pathways was investigated using NanoString PanCancer Immune Profiling Panel.

**Results:** High-MAC and high-MACi diets increased fecal microbial alpha-diversity and the abundances of SCFA producing families *Lachnospiraceae,* and *Oscillospiraceae* as well as fecal SCFA levels, compared with low-MAC diet. E0771 tumors responded to anti-PD1 in mice fed high-MAC, while mice fed high-MACi did not respond. Low-MAC fed mice with single E0771 allograft also responded to anti-PD1, but genistein supplementation eliminated responsiveness. E0771 tumors in high-MAC fed mice contained elevated levels of exhausted CD8+ T cells, which were decreased after anti-PD1 therapy. Opposite effects were seen in mice fed high-MACi diet. Mice with DMBA-initiated ERα+ mammary tumors did not respond to anti-PD1. TAM converted TNBC and ERα+ tumors to become sensitive to anti-PD1 therapy in mice fed high-MACi or low-MAC diets, respectively. Genes in TH17 differentiation pathways were linked to TAM-induced improved anti-PD1 response both in TNBC and ERα+ mammary tumors.

**Conclusions:** Our results highlight the role of diet in impacting the effectiveness of ICB therapies. We found that increased SCFA levels alone are not predictive of response to anti-PD1, but if tumor expresses ERα or if diet contains ERα activating compounds, such as isoflavones, blocking ERα+ might convert unresponsive tumors responsive to anti-PD1. **Word count**: 339

**What is already known on this topic:** Dietary fiber is proposed to improve response to checkpoint inhibitor therapy against melanoma, but this has been challenged by a recent preclinical study in which different mouse tumor models were used. None of the studies have been done in breast cancer or preclinical breast cancer models.

**What this study adds:** Our study showed, using triple negative breast cancer (TNBC) and estrogen receptor positive (ER+) breast cancer models, that indeed high levels of microbiota accessible carbohydrates (MACs) in diet did not alone determine responsiveness to anti-PD1 therapy, but diet high in plant isoflavones/ hormones impaired anti-PD1 effectiveness, regardless of whether diet contained high fiber levels or not. We also found that the adverse effects of isoflavones were counteracted by tamoxifen, partial estrogen receptor antagonist.

**How this study might affect research, practice or policy:** Our findings could indicate that breast cancer patients, both those with TNBC and ER+ disease, should not consume diets high in isoflavones when treated with anti-PD1.

## Introduction

Factors determining effectiveness of cancer immunotherapies, such as those targeting programmed cell death protein 1 (PD1) and its ligand PDL1, remain elusive. Diet might influence responsiveness: observational evidence, confirmed in animal studies, suggests that melanoma patients consuming high levels of dietary fibers are most responsive to immunotherapies while prebiotics may have an opposite effect (1). Beneficial effects of fiber on immune responses, including PD1/PDL1 therapies, are thought to be mediated by a fiber-induced increase in the abundance of gut bacteria that produce short-chain fatty acids (SCFAs) (2–4). However, the beneficial role of dietary fiber in anti-PD1 or anti-PDL1 therapy has been recently challenged by a study involving diverse murine tumor models (5). Breast cancer was not among the models.

Microbiota-accessible carbohydrates (MACs) are a group of fermentable dietary fibers (6) that increase gut bacterial production of SCFAs (7). However, many high-MAC diets that are fed to laboratory mice contain estrogenic isoflavones which might modify immune therapy response (8). Human dietary sources of MACs, such as vegetables, legumes, fruits and berries, also contain high levels of isoflavones. It has been reported that activated estrogen receptor α (ERα) promotes immunosuppression (9, 10). Estrogens also drive gender-dependent immunosuppression in the tumor microenvironment (TME) (11), including driving differentiation of naïve CD4+ T cells into immunosuppressive Treg cells (12): these effects are purported to explain why men might respond better to cancer immunotherapies than women (13). Further, inhibiting ERα with fulvestrant improved response to immunotherapies against estrogen-insensitive, preclinical, non-breast tumor models (9, 14).

We investigated here whether feeding mice high– or low-MAC diets, or a high-MAC diet containing isoflavones from soymeal (MACi), affected responsiveness to anti-PD1 therapy against triple negative breast cancer (TNBC) or ERα+ mammary tumors in female mice. In addition, the impact of blocking ERα+ with tamoxifen (TAM) on ICB effectiveness was studied. Our results indicated that high-MAC and MACi diets increased fecal SCFA producing bacteria and SCFA levels. However, high fecal SCFA levels alone did not predict responsiveness to anti-PD1, since anti-PD1 reduced TNBC growth in mice fed high– or low-MAC diet but not in mice fed high-MACi diet. The latter observation suggests that isoflavones impaired ICBs effect even in mice that had high fecal SCFA levels. TAM converted TNBCs in high-MACi fed mice towards responsiveness, while ERα+ mammary tumors started responding to anti-PD1 if mice were fed low-MAC diet and treated with TAM. Thus, TAM’s effect was dependent on tumor hormone receptor status, and the MAC diet mice were consuming.

## Materials and Methods

### Mice

Four-week-old, female C57BL/6 mice were obtained from Taconic Biosciences, Inc. The studies were performed at Georgetown University and at University of Minnesota, keeping mice at 22±2°C with *ad libitum* access to diet and chlorinated filtered drinking water, and approved by their respective Institutional Animal Care and Use Committees. At Georgetown University, mice were maintained on a 12-h light-dark cycle in a specific-pathogen-free facility on HEPA-filtered ventilated racks, while at University of Minnesota, mice were housed in a conventional facility in static cages with filter tops on a 14:10 light/dark cycle. Mice were euthanized at the end of the study using carbon dioxide inhalation and tissues harvested as needed.

### Diets

For studies 1 and 2, mice were fed irradiated diets that were either low-MAC with no isoflavones (LabDiet AIN93G, Land O’Lakes Inc.), high-MAC containing low levels of isoflavones (LabDiet 5V5M), or high-MAC containing moderate levels of isoflavones (LabDiet 5058D). For study 3, mice were fed low-MAC diet (AIN93G; TD.97184) or low-MAC diet supplemented with 500 ppm genistein (AIN93G+500 ppm genistein; TD.210530) from Envigo Teklad Diets. According to Envigo Teklad Diets, high-MAC isoflavone (high-MACi) diet contained 125-275 ppm of these isoflavones. Basic nutritional comparisons of these diets are provided in **Supplementary Table 1**. While the diets were not nutritionally identical, the percentage and source of protein, fat and carbohydrate in the three diets were roughly similar.

### Gut microbiome analysis: Amplicon sequencing and bioinformatic processing

Fecal pellets were collected from mice fed low-MAC, high-MAC or high-MACi diet for four weeks. These mice did not have any mammary tumors. The V4 hypervariable region of the 16S rRNA gene was amplified and sequenced using the 515F/806R primer set by the University of Minnesota Genomics Center (UMGC) (15). Raw sequence data has been deposited in the NCBI SRA under BioProject accession number <u>SRP579135</u>. Sequence data was processed using mothur (ver. 1.41.1) (16) and a modified version of our previously published pipeline (17).

### Assessment of fecal SCFA levels

For SCFA analysis, fecal samples were collected and snap-frozen from mice fed low-MAC, high-MAC, or high-MACi diets. To obtain sufficient material, feces from 2–3 mice were pooled, yielding three biological replicates per group. All SCFA analytical standards were purchased from Sigma-Aldrich (MO, USA). Fecal SCFA analysis was performed using a gas chromatography-coupled mass spectrometry (GC-MS) platform as previously described (18). Quantitation of SCFAs from the raw GC-MS data was performed using a 7-point calibration curve with appropriate solvent blanks, negative controls, and quality control samples run throughout the assessment and by utilizing the open-source software Skyline (19). Calibration curves were fitted independently for each compound by linear in log space regression using the peak ratio of each compound to the global internal standard. All calibration curves were fit with an R2 of at least 0.995 precision. All estimated sample concentrations were normalized by starting lyophilized material weight.

### Effect of low– and high-MAC diets and tamoxifen (TAM) on anti-PD1 response in TNBC model

We investigated whether three laboratory diets affected the growth of ERα-negative murine E0771 mammary tumors in syngeneic C57BL/6NTac mice. After acclimation, mice were assigned to low-MAC (n=33), high-MAC (n=33), or high-MACi (n=32) diets. At 8 weeks of age, after 4 weeks on diet, mice were allografted with 1×10⁶ E0771 cells in medium:Matrigel (1:1) into both 4th mammary fat pads. When tumors reached 50–70 mm³, mice were subdivided into four treatment groups (8–9 mice): IgG control, tamoxifen (TAM) pellet (5 mg, 60-day release), anti-PD-1 mAb (125 µg IP; clone RMP1-14, BioXcell), or TAM+anti-PD-1. Anti-PD-1 was administered in three doses at 3–4-day intervals; controls received a sham pellet and rat IgG (clone 2A3, BioXcell) on the same schedule.

To account for enhanced immune activation from dual-tumor allografting (20, 21), a second experiment evaluated diet effects in mice bearing a single tumor. Here, 0.5×10⁶ E0771 cells in PBS:Matrigel (1:1) were implanted into one mammary fat pad (9–10 mice/group). Upon reaching 50–70 mm³, mice received either 3 doses of 50 µg anti-PD-1 or IgG2a control; no hormone therapy was used. Tumors in both studies were measured twice weekly, volumes calculated as ½ × (length × width²), and mice were monitored for up to 4 weeks after the final treatment or euthanized earlier if tumor burden exceeded 1500 mm³.

### Effect of genistein on anti-PD1 response

C57BL/6NTac mice were assigned to either a low-MAC diet or the same diet supplemented with 500 ppm genistein (genistein group; genistein was prepared in Dr. Helferich’s lab) for 4 weeks. Mice were allografted with 0.5 × 10^6^ E0771 cells in 1× PBS mixed with Matrigel (1:1) into the right 4th mammary fat pad. Once tumors reached 50-70mm^3^, mice (n=10/group) were treated with either control IgG mAb or anti-PD1 mAb. This experiment was repeated twice. In the first study, mice were treated with 150 µg anti-PD1 mAb or control IgG, and in the replicate, a reduced 100 µg anti-PD1 or IgG was used.

### Effects of low– and high-MAC diets on anti-PD1 therapy in ERα+ mammary tumor model

This study evaluated whether DMBA-induced ERα⁺ mouse mammary tumors respond to anti-PD1 therapy and whether adding TAM modifies this response. Unlike rats, mice with ERα⁺ DMBA-induced tumors do not exhibit tumor growth reduction with TAM (22). Female C57BL/6NTac mice were fed low– or high-MAC diets from age 4 weeks and received medroxyprogesterone acetate (MPA) at week 6, followed by 1 mg DMBA doses at weeks 7–10, generating nearly 100% tumor incidence. Once tumors reached 50–70 mm³ (∼12.6 weeks after the last DMBA dose), mice were divided into four treatment groups: (1) 200 μg IgG vehicle and placebo pellet, (2) 200 μg of anti-PD1, (3) TAM ( 5 mg as a 60-day release pellet) or (4) a combination of anti-PD1 and TAM. Each group contained 7-11 mice. Higher anti-PD1 doses were used anticipating a weak response in this ERα⁺ mammary tumor model. Tumor growth was monitored twice weekly over 11 weeks.

### Immune cell analysis in E0771 mammary tumor model

Tumors (n=4-6/group) were harvested 12 days after last anti-PD1 dose to assess the effect of diets on tumor-infiltrating immune cells. Tumor growth data in these mice is shown in Fig. 4A-C. Single-cell suspensions were prepared by mechanical dissociation using 70 µm sterile nylon strainers and resuspended in T cell media. Approximately 2 x 10^6^ cells were stimulated for 3 hours at 37°C using 50 ng/ml of Phorbol 12-myristate 13-acetate (Sigma-Aldrich), 750 ng/ml of Ionomycin (Sigma-Aldrich) and 10 μg/ml of brefeldin (Biolegend). Cells were stained for viability using the LIVE/DEAD Fixable Near IR Dead cell kit (Invitrogen), fixed with 4.2% formaldehyde buffer, and permeabilized with Perm/Wash buffer (BD Biosciences). Immunophenotyping was performed using conjugated antibodies against CD3 (clone: 145-2C11), CD8 (clone: 53-6.7), and TIM3 (clone: B8.2C12). Data were acquired using LSRFortessa Flow Cytometer and analyzed by Flowjo 10.9-10. Positive staining gates were defined using Fluorescence Minus One (FMO) control, and compensation was performed using the AbC™ Total Antibody Compensation Bead Kit (Invitrogen).

### Expression of ERα and ERβ in breast cancer cells

Murine E0771, and human MCF-7 and T47D breast cancer cells were cultured in RPMI 1640 medium supplemented with 5% FBS. Once MCF-7 and T47D cells reached 70% confluence, the medium was replaced with phenol red-free RPMI 1640 supplemented with 5% charcoal-stripped FBS for five days. Following hormone deprivation, the cells were treated with 10 nM 17β-estradiol (E2) for 24 hours. E2 was diluted in ethanol, with the final ethanol concentration kept below 0.1%.

Protein extraction and Western blotting were conducted as previously described (23), with non-specific binding blocked using 4% Bovine Serum Albumin and primary antibodies against ERα (Abcam,ab16460 and ERβ (Novusbio, NB120-3577) used at a $1:1000 dilution.

### Effect of isoflavone genistein on TNBC growth in *vitro*

E0771 cells (1 × 10³) were seeded in a 96-well plate and, after 24 hours, treated with genistein (5 μM), tamoxifen (1 μM), a combination of genistein and tamoxifen, or vehicle control (0.04% Dimethyl sulfoxide-DMSO). Independent experiments (n=4) were performed with three technical replicates. Cell growth was monitored using the Incucyte SX-5 system (Sartorius, Germany) for up to 120 hours, with confluence analyzed as a percentage relative to the initial measurement at 0 hours.

### Nanostring analysis

The PanCancer Immune Profiling Panel containing 770 genes, including 40 PanCancer reference genes, purchased from NanoString Technologies, was used to assess possible differences in immune signaling pathways among mice fed high MACi diet and treated with IgG alone, anti-PD1 alone, TAM alone or the combination of both anti-PD1 and TAM. For this analysis, we used three tumors from three mice per group, harvested at the end of the tumor monitoring period.

### Gene expression by PCR

For confirmation of Nanostring data, we performed qRT-PCR analysis in mammary tumors (23) Primers used in qRT-PCR analysis were designed using the IDT tool (Integrated DNA Technologies Coralville, IA, USA, primer sequence found in **Supplementary Table 2**).

### Data Analysis

#### Microbiome data analysis

Alpha diversity was calculated as the Shannon index using mothur. Beta diversity was calculated using Bray-Curtis dissimilarity matrices and visualized via ordination using principal coordinate analysis (PCoA). Differences in alpha diversity indices and relative abundances of taxa were determined using Kruskal-Wallis test in XLSTAT (ver. 2020.5.1; Addinsoft, New York). Differences in community composition were determined using analysis of similarity (ANOSIM) (24), and taxa that were significantly correlated with either PCoA axis by Spearman correlation, using the corr.axes function in mothur, were overlaid on the plot.

#### Nanostring data analysis

We used the ruv package (v0.9.7.1) in R (v4.1) to normalize the raw count and the batch effect for Nanostring data. Pair wise comparison among experimental groups were conducted using limma package (v3.58.1). Differentially expressed genes with fdr < 0.05 were selected and used as inputs for Gene Set Enrichment Analysis (GSEA) (v4.2.3, Broad Institute), with KEGG gene set used for enrichment score calculations.

#### Data analysis for all other end-points

Differences in SCFA and BCFA levels among groups were investigated by one-Way ANOVA followed by Tukey’s multiple comparisons test. Differences in tumor growth/burden among experimental groups were analyzed by two-way repeated measures ANOVA followed by Student-Newman-Keuls Post-Hoc test. Differences in E0771 mammary tumor “take” in mice allografted cells in both 4^th^ mammary glands were analyzed by Chi2-test. The effects of genistein and tamoxifen treatment on E0771 cells *in vitro* were analyzed by two-way repeated measures ANOVA. Differences in tumor multiplicity, percentage of CD8+ and CD8+TIM3+ T cells in the tumors, and expression of genes in Th17 pathway were analyzed by two-way ANOVA followed by Tukey’s multiple comparisons test. GraphPad Prism and R package agricolae (v1.3-7) were used to perform these statistical analyses. Results with p values <0.05 were considered statistically significant.

## Results

### High– and low-MAC diets have differential effects on the gut microbiome

Alpha diversity was significantly greater in mice fed high-MAC or high-MACi diet, compared with low-MAC diet (**Fig. 1A**). Communities were predominantly comprised of members of the SCFA producing families *Muribaculaceae*, *Lachnospiraceae*, and *Oscillospiraceae* (**Table 1**; **Fig. 1B**). Like alpha diversity, significant differences among relative abundances of most taxa were seen between low– and the two high-MAC diets, with little distinction between high-MAC and high-MACi diets (**Table 1**). *Muribaculum* of *Muribaculaceae* family was the only genus that had significantly greater relative abundance, roughly twofold, in the high-MAC diet relative to both high-MACi and low-MAC diets. However, *Muribaculaceae* family was significantly higher in low-MAC than the two high-MAC diets. According to these findings, significantly higher abundances of *Muribacululaceae* family and its *Muribaculum* genus was seen in mice that were responsive to anti-PD1. Community composition structure, as evaluated using Bray-Curtis dissimilarities, was significantly different among all groups (ANOSIM R = 0.53, *P* < 0.001; **Fig. 1C**), which was supported by post-hoc pairwise comparisons (R = 0.27 – 0.71, *P* ≤ 0.004, at Bonferroni-corrected *α* = 0.017).

**Figure 1.**
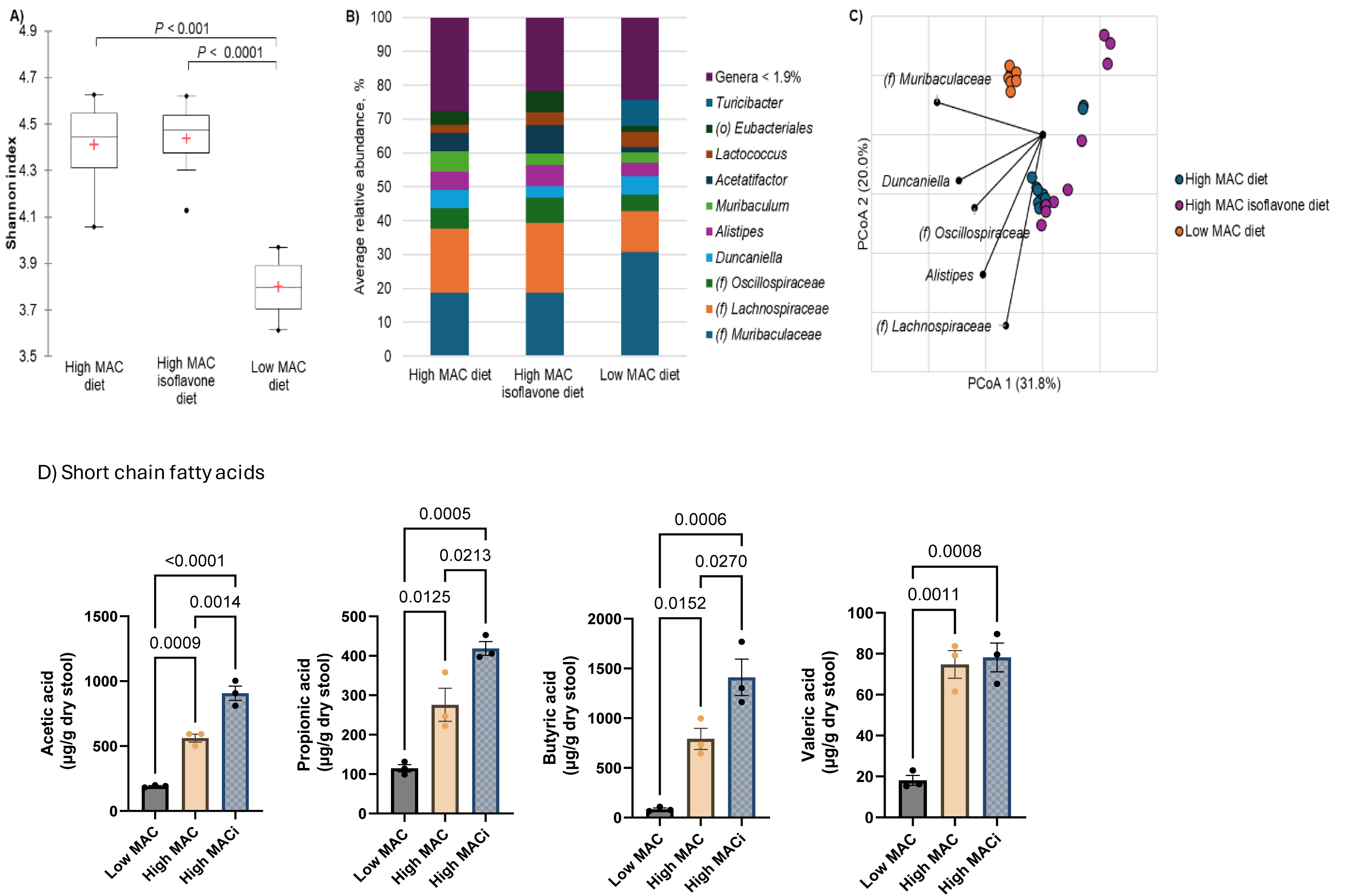
Microbiome diversity and composition and short-chain fatty acid (SCFA) levels in fecal samples of mice receiving different diets. In mice fed low-MAC, high-MAC or high-MAC isoflavone (MACi) diet (A) Box plots of Shannon diversity; (B) distribution of predominant genera classified to genus or most resolved taxonomic level. (f) and (o) indicate taxa that could not be classified passed family or order, respectively; (C) Principal coordinate analysis of Bray-Curtis dissimilarities (r^2^ = 0.82); amd (D) Fecal SCFA levels. C57BL/6Tac mice were kept on one of the three diets for 4 weeks, after which fecal pellets were collected from 10 mice per group. Fecal pellets for SCFA assays were pooled to have three biological replicates per group. Data are shown as mean ± SEM with statistical significances obtained by one-way ANOVA.

**Table 1.** Distribution of predominant taxa under different diets.

| **Taxa** | **High-MAC diet** | **High-MACi diet** | **Low-MAC diet** | **P-value** |
| --- | --- | --- | --- | --- |
| *(o) Eubacteriales* | 4.16 ± 0.96 A | 6.19 ± 2.58 A | 1.71 ± 0.31 B | < 0.0001 |
| *(f) Lachnospiraceae*<br>(of *Eubacteriales*<br>order) | 18.87 ± 3.47 A | 20.57 ± 5.71 A | 12.15 ± 2.58 B | <0.001 |
| *(f) Oscillospiraceae*<br>(of *Eubacteriales*<br>order) | 6.04 ± 0.78 | 7.44 ± 3.59 | 4.88 ± 0.86 | 0.059 |
| *(f) Muribaculaceae* | 18.76 ± 5.81 B | 18.75 ± 2.98 B | 30.70 ± 4.16 A | <0.001 |
| *Muribaculum* (of<br>*Muribaculaceae*<br>family) | 6.10 ± 2.40 A | 3.45 ± 1.70 B | 2.99 ± 0.37 B | 0.001 |
| *Duncaniella* (of<br>*Muribaculaceae*<br>family) | 5.48 ± 2.49 | 3.60 ± 1.63 | 5.36 ± 2.43 | 0.152 |
| *Alistipes* | 5.34 ± 1.48 AB | 6.18 ± 1.47 A | 4.02 ± 1.28 B | 0.016 |
| *Acetatifactor* | 5.38 ± 2.72 A | 8.26 ± 3.02 A | 1.60 ± 0.32 B | <0.001 |
| *Lactococcus* | 2.32 ± 1.89 A | 3.70 ± 1.58 A | 4.50 ± 1.82 A | 0.048 |
| *Turicibacter* | 0.01 ± 0.01 B | 0.01 ± 0.01 B | 7.64 ± 4.22 A | < 0.0001 |

Taxa are classified to genus or most resolved taxonomic level. (f) and (o) indicate designations could only be made to family and order, respectively.

### High-MAC diets increased fecal SCFA levels

Both high-MAC diets (high-MAC and high-MACi) significantly increased the concentrations of acetic acid (p=0.009 and <0.0001), propionic acid (p=0.01 and p=0.0005), butyric acid (p=0.015 and p=0.0006) and valeric acid (p=0.0011 and p=0.0008) in stool, compared with low-MAC diet (**Fig. 1D**). The levels of acetic acid (p=0.001), propionic acid (p=0.02) and butyric acid (p=0.02) were significantly higher in high-MACi than high-MAC fed mice (**Fig. 1D**).

### Study 1

#### TNBC model

##### Effect of low– and high-MAC diets on TNBC growth and effectiveness of anti-PD1 treatment

Fifty-two percent (52%) of the E0771 tumors allografted to the 4^th^ mammary glands of both the right and left side of a mouse that were fed low-MAC diet did not grow in either mammary gland, i.e., tumor cells might have been cleared by the immune system (**Fig. 2A**). Although tumors failed to grow in 15% of the allografted tumors in high-MAC fed mice and in 17% of the tumors allografted to high-MACi fed mice, these values were significantly lower than 52% in low-MAC diet fed mice (p<0.0001; Chi-square test). If one tumor did not start to grow, the other in the same mouse did not grow either. Further, two mammary tumors within the same mouse showed the same trend in tumor progression or regression, i.e., both “ tumor-take” and the response to anti-PD1 was mouse centric rather than tumor centric, as also reported by others (25).

**Figure 2.**
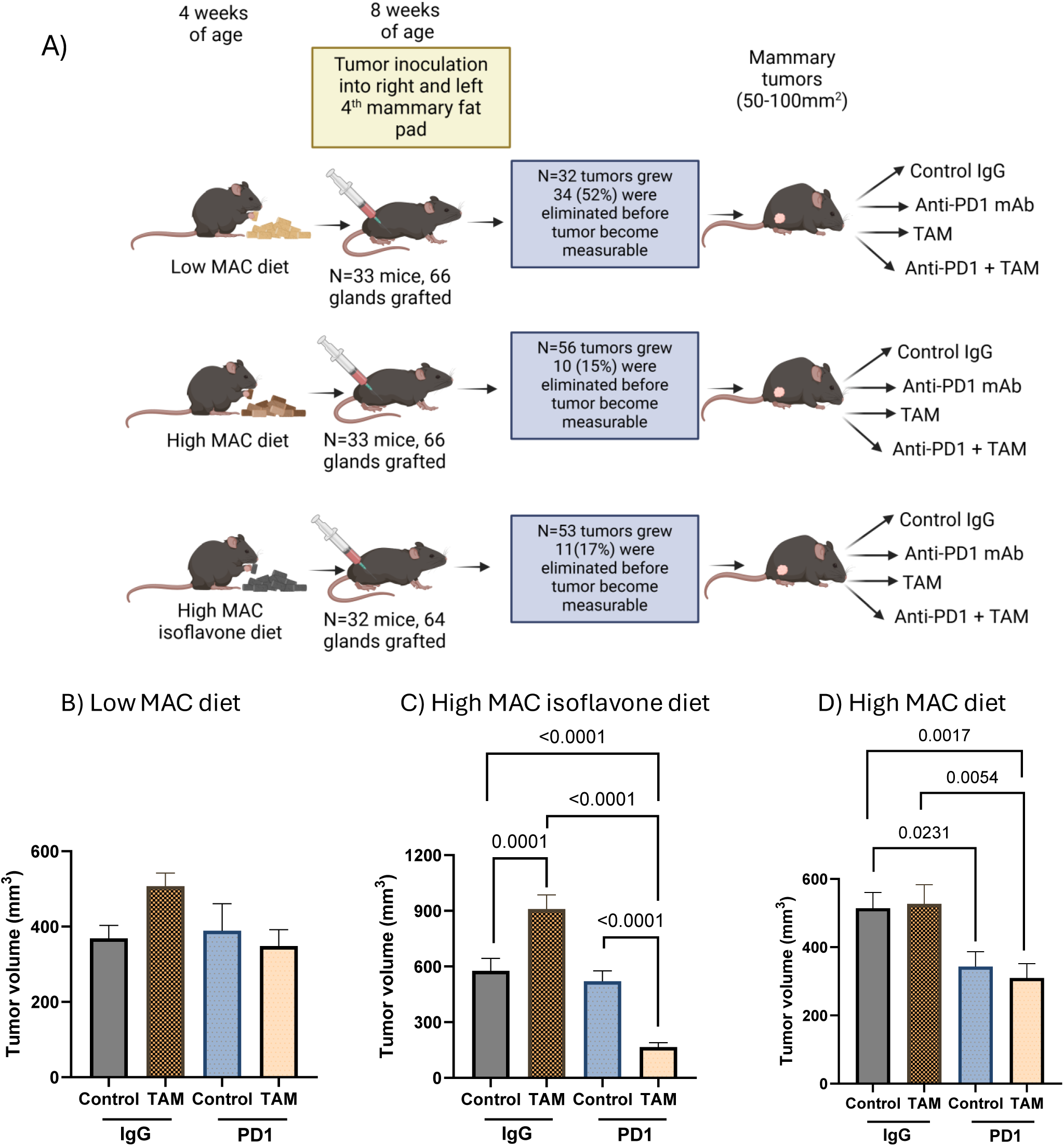
Effect of anti-PD1 and/or tamoxifen on the growth of E0771 TNBC in mice fed different diets. (A) Experimental design, (B,C,D) tumor burden in mice treated with IgG, tamoxifen (TAM), anti-PD1 or TAM+anti-PD1 and fed (B) low-MAC diet, (C) high-MAC diet, or (D) high-MAC isoflavone diet (MACi). Each mouse was allografted with two tumors: one on the right and another on the left 4^th^ mammary gland. Tumor burden was calculated by combining the volume of right and left tumors and then as an average of the measurenment period per mouse, according to R package agricolae (v1.3-7). There were 4-8 mice in each group. Data are shown as mean ± SEM with statistical significances obtained by two-way repeated measures ANOVA followed by Student-Newman-Keuls Post-Hoc test. Experimental design illustration was created with BioRender.com

Anti-PD1 treatment had no effect on mammary tumor growth in animals fed either a low-MAC (**Fig. 2B**) or high-MACi diet (**Fig. 2C)**. In the high-MAC group, a significant inhibition in tumor growth was observed by anti-PD1 therapy, compared with IgG controls (p=0.023) (**Fig. 3D**).

**Figure 3.**
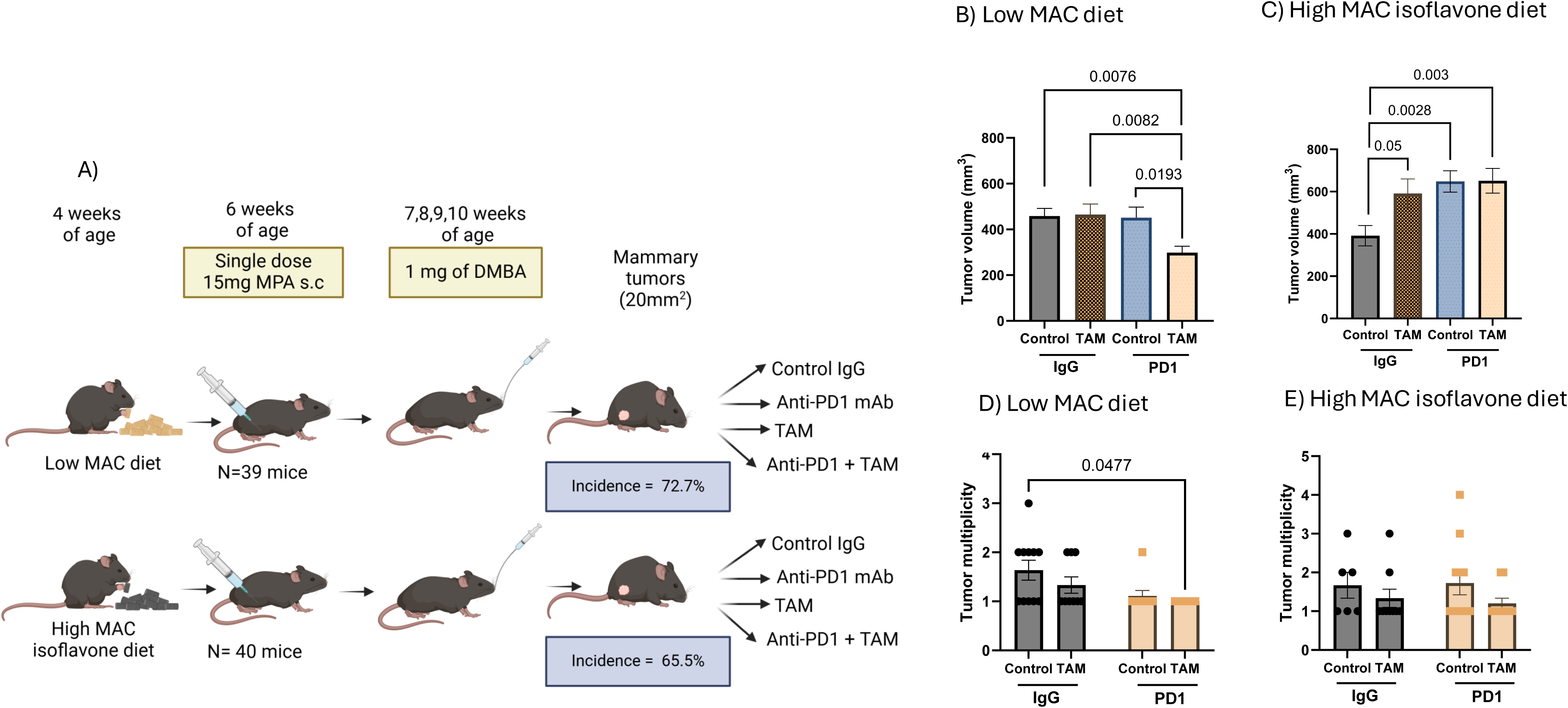
Effect of anti-PD1 and/or tamoxifen on the growth of ERα+ tumors in mice fed different diets. (A) Experimental design of 7,12-dimethylbenz[a] anthracene (DMBA)-initiated estrogen receptor α positive (ERα+) mammary tumors in mice fed low-MAC or high-MAC isoflavone (MACi) diet, (B,C) tumor burden in mice treated with IgG, TAM, anti-PD1 or TAM+anti-PD1 and fed (B) low-MAC diet, or (C) high-MACi diet. Tumor burden per mouse was calculated by combining the volume of all tumors and then as an average of the measurenment period per mouse, according to R package agricolae. Tumor multiplicity in mice treated with IgG, TAM, anti-PD1 or TAM+anti-PD1 and fed (D) low MAC diet, or (E) high MACi diet. There were 7-11 mice in each group. Data are shown as mean ± SEM with statistical significances obtained by two-way repeated measures ANOVA followed by Student-Newman-Keuls Post-Hoc test for tumor burden and two-way ANOVA followed by Tukey’s multiple comparisons test for tumor multiplicity. Experimental design illustration was created with BioRender.com

**Figure 4.**
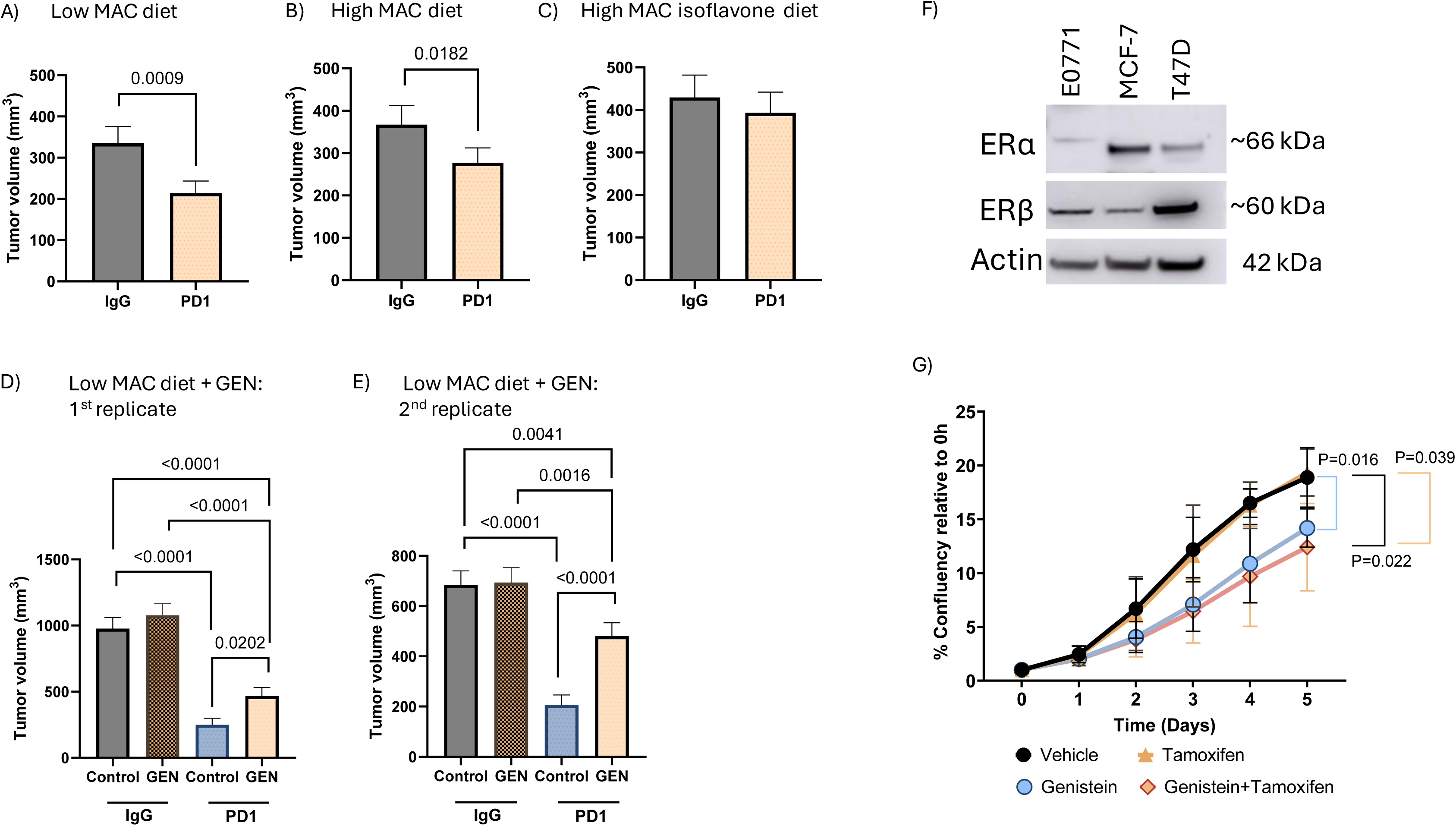
Effect of genistein (GEN) on anti-PD1 response *in vivo*, and the growth of E0771 TNBC cells *in vitro*. Mice were fed (A) low-MAC, (B) high-MAC, (C) high-MAC isoflavone (MACi) diet, or (D,E) low-MAC diet supplemented with 500 ppm GEN. Each mouse was allografted with one tumor on the right 4^th^ mammary gland. Tumor burden was calculated as an average of the measurenment period per mouse, according to R package agricolae. There were 7-10 mice in each group. Data are shown as mean ± SEM with statistical significances obtained by two-way repeated measures ANOVA followed by Student-Newman-Keuls Post-Hoc test. (F) Protein levels of estrogen receptor α (ERα) and ERβ in E0771 murine mammary tumor cells and human ER+ MCF-7 and T47D breast cancer cell lines. (G) Growth of E0771 tumor cells treated with genistein (GEN: 5 μM), tamoxifen (TAM: 1 μM), a combination of genistein and tamoxifen, or vehicle control (0.04% DMSO) for up to 120 hours. Data are shown as mean ± SEM of four independent experiments with three technical replicates each. Statistical significances shown were obtained by two-way repeated measures ANOVA followed by Student-Newman-Keuls Post-Hoc test.

##### Effect of anti-estrogen tamoxifen (TAM) on E0771 mammary tumor growth in mice fed low– or high-MAC diets

Since isoflavones bind and activate ERα (26–28), and activation of ERα activates immunosuppressive cells (9, 10), we questioned whether the lack of efficacy of anti-PD1 therapy in mice fed high-MACi diet might be due to the isoflavone genistein activating ERα in the TME. We explored this by treating mice with TAM, a partial ERα antagonist. TAM has been reported to block ERα in the tumor and increase infiltration of CD4+ T cells (29) and activate CD8+ T cells (30) in the TME. In breast cancer patients, TAM suppressed MDSCs (31). Estradiol is reported to have an opposite effect on MDSCs (9, 10). In this study, mice were allografted E0771 mammary tumors to both left and right 4^th^ mammary glands. Among low-MAC and high-MAC diet fed mice, tumor burden was similar in IgG control and TAM treated mice (**Fig. 2B,D**). Unexpectedly, mice fed high-MACi diet exhibited increased E0771 tumor growth when treated with TAM, compared with IgG treated mice (p=0.0001) (**Fig. 2C**).

##### Effect of diet, tamoxifen and anti-PD1 treatment on tumor growth

Treatment with a combination of anti-PD1 plus TAM significantly inhibited mammary tumor growth in mice fed high-MACi diet, compared with TAM or anti-PD1 treatments as monotherapies or with IgG control (p<0.0001) (**Fig. 3C**). The combination treatment did not further reduce tumor growth in high-MAC-diet-fed mice, compared with anti-PD1 treatment alone (**Fig. 3D**), nor did it affect tumor growth in low-MAC-diet-fed mice (**Fig. 3B)**. Thus, although TAM on its own promoted tumor growth in high-MACi group, in combination with anti-PD1 it induced highly significant tumor growth reduction. These data suggest that blocking ERα in the TME of TNBC is required for anti-PD1 to be effective in an elevated estrogenic environment.

#### ERα+ breast cancer model

##### Effect of low– and high-MAC diets on ERα+ mammary tumor growth and anti-PD1 response

We compared the effect of low-MAC and high-MACi diets on anti-PD1 responsiveness in ERα+ DMBA model. Experimental design of the study is shown in **Fig. 3A**.

The incidence of DMBA-induced tumors was similar in the two dietary groups (**Fig. 3A**). Among mice that were fed high-MACi, 65.5% of mice developed mammary tumors (multiplicity 1.5 ± 0.13 per mouse), and among mice fed low-MAC diet, 72.7% of mice developed mammary tumors (multiplicity 1.3 ± 0.09 per mouse). Thus, diet did not affect the incidence or multiplicity of mammary tumors. In low-MAC diet group, anti-PD1 did not impact tumor growth (**Fig. 3B**). When compared with IgG controls, anti-PD1 therapy in mice fed high-MACi diet unexpectedly significantly promoted tumor growth (p=0.0028) (**Fig. 3C**).

##### Effect of diet and tamoxifen on tumor growth

TAM stimulated DMBA-induced mammary tumor growth in mice fed a high-MACi diet (p=0.05) (**Fig. 3C**) but did not affect tumor growth in mice fed a low-MAC diet (**Fig. 3B**). These results are like those observed in mice allografted with E0771 TNBC, i.e., TAM increased tumor growth when given with isoflavone containing diet, regardless of whether the tumors expressed ERα. These findings suggest that the tumor growth potentiating effects of TAM in mice fed high-MACi diet were not linked to ERα status of the tumors, but the effects of isoflavones on the TME.

##### Effect of diet, tamoxifen and anti-PD1 treatment on tumor growth

The combination treatment with TAM and anti-PD1 inhibited tumor growth in low-MAC fed mice with ERα+ mammary tumors compared with IgG (p=0.0076), or anti-PD1 (p=0.0193) or TAM (p=0.0082) monotherapy treated mice (**Fig. 3B**). The combination treatment also decreased tumor multiplicity compared to mice treated with IgG (p=0.048) in mice fed low-MAC diet (**Fig.3D**). In contrast, in the high-MACi group, tumor growth during combination treatment was like that seen in mice treated with TAM or anti-PD1 alone (**Fig. 3C)**. There was no difference in multiplicity among mice fed high-MACi diet **(Fig. 3E)**. These findings are opposite to the results obtained in mammary tumors from mice allografted with E0771 cells. In the TNBC model, TAM+anti-PD1 effectively reduced E0771 tumor growth when mice were fed high-MACi diet. Thus, the presence or absence of ERα in mammary tumors might impact how diet and TAM modify response to anti-PD1 therapy.

### Study 2

#### Allografting E0771 cells to a single mammary gland per mouse

In this experiment, each mouse was allografted E0771 tumor cells into the right 4^th^ mammary gland only, resulting 100% of allografted tumors to start to grow in low-MAC, high-MAC and high-MACi fed diets. Anti-PD1 was now effective in inhibiting tumor growth both in mice that were fed low-MAC (p=0.0009) (**Fig. 4A**) and high-MAC diet (p=0.0182) (**Fig. 4B**). However, like in Study 1 (**Fig. 2C**), mice fed high-MACi diet did not respond to anti-PD1 therapy (**Fig. 4C**).

### Study 3

#### Effect of the isoflavone genistein on anti-PD1 response

Mice with a single E0771 tumor allograft, fed low-MAC diet anti-PD1 at doses of 100 μg or 150 μg significantly reduced tumor burden compared with control IgG (p<0.0001) (**Fig. 4D,E**). Adding genistein to low-MAC diet significantly reduced the effect of anti-PD1 on tumor growth (p=0.0202 and p<0.0001 between anti-PD1-control and anti-PD1-genistein mice for 1^st^ replicate and 2^nd^ replicate, respectively) (**Fig. 4D,E**).

### Effect of tamoxifen on genistein treated E0771 mammary tumor cells *in vitro*

E0771 mammary tumor cells did not express ERα protein but were positive for ERβ (**Fig. 4A)**. Consistent with these cells being ERα negative, 1 μM TAM did not affect their growth (**Fig. 4B**). However, 5 µM genistein significantly inhibited E0771 tumor cell proliferation (p=0.016) (**Fig. 4B**). The combination of TAM + genistein did not further inhibit growth, i.e., TAM did not add to the growth inhibitory effect of genistein. Since genistein binds to cancer growth suppressing ERβ (32), these findings suggest that activation of ERβ by genistein in tumor cells, in the absence of ERα, can directly inhibit the growth of E0771 mammary tumors. The data also indicate that TAM’s E0771 tumor growth promoting effect *in vivo* in high-MACi-fed mice likely was caused by the combined effect of TAM and isoflavones on the TME.

### Effect of low– and high-MAC diets on tumor CD8+ T cell infiltration and exhaustion in anti-PD1 treated mice

High-MAC-diet-fed mice exhibited a significantly higher proportion of exhausted TIM3+ CD8+ cells in the TME than mice fed a low-MAC diet (p=0.017) or high-MACi diet fed mice (p=0.022) (**Fig. 5A**). Infiltration of CD8+ T cells was similar among the three dietary groups (**Fig**. 5**B**). Treatment with anti-PD1 significantly suppressed the TIM3+ CD8+ T cell population in the TME of high-MAC diet fed mice (p=0.013) but increased these cells in high-MACi group (p=0.002 for interaction between diet and anti-PD1). However, anti-PD1 increased CD8+ T cell infiltration in low-MAC diet group (p=0.03) (**Fig. 5B**), indicating a stronger anti-tumor immune response. The findings obtained here are consistent with mice fed high-MAC diet being responsive to anti-PD1 therapy, while high-MACi diet fed mice being resistant.

**Figure 5.**
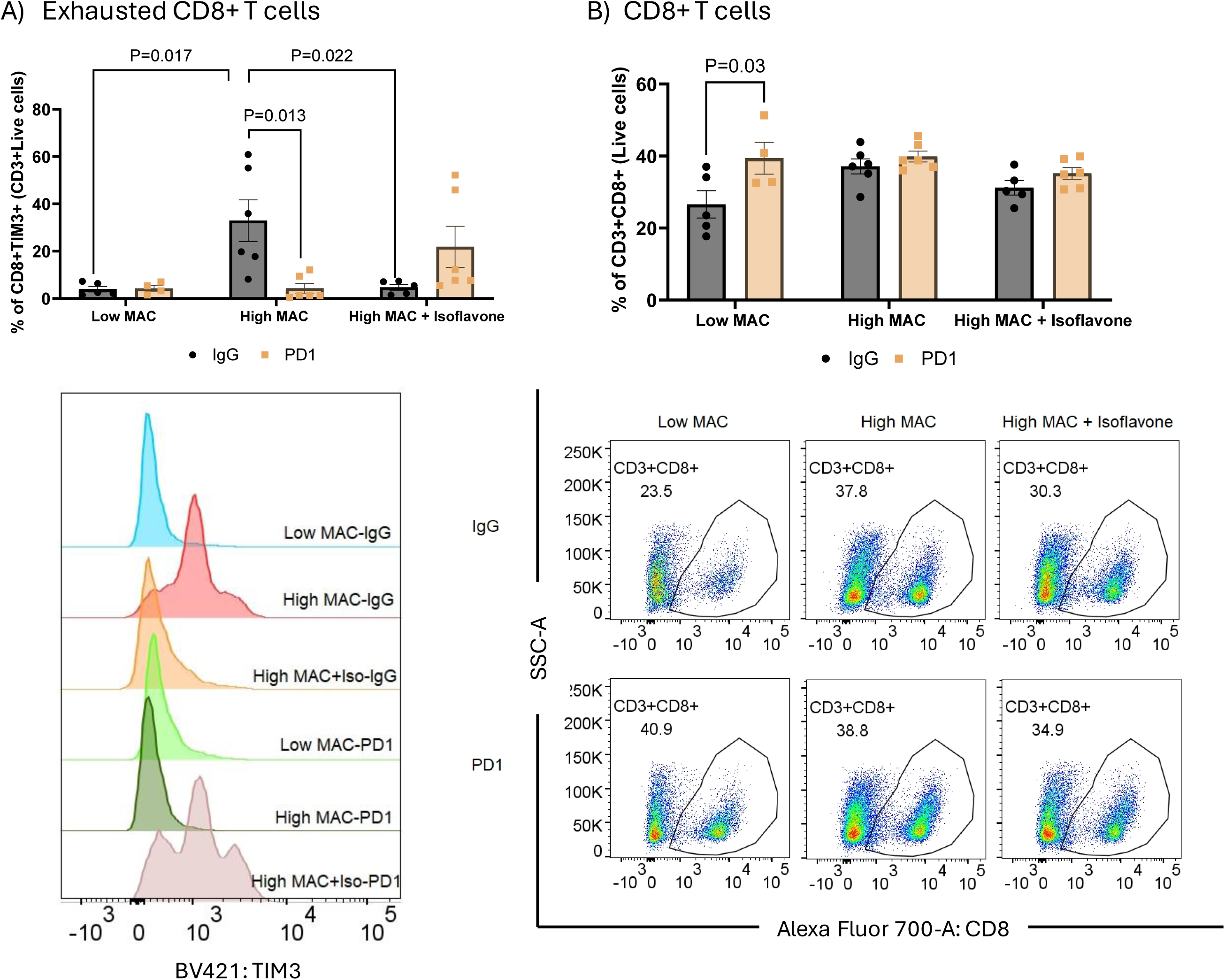
Effect of anti-PD1 on CD8+ T cell exhaustion in mice fed different diets. (A) Frequency of CD8+ exhausted cells (CD8+TIM3+) and (B) total CD8+ cells (CD3+CD8+) in tumors of mice fed low-MAC, high-MAC or high-MAC isoflavone (MACi) diets and treated with anti-PD1 or anti-IgG. Cells were gated on live cells, followed by CD3+ cells, CD8+ cells and TIM3+ cells. Gates were determined based on FMO staining. There were 4-6 tumors in each group. Data are shown as mean ± SEM with statistical significances obtained by two-way ANOVA followed by Tukey’s multiple comparisons test.

### NanoString analysis in high-MACi diet fed mice

Since TAM sensitized E0771 tumors in mice fed high-MACi diet to anti-PD1, a NanoString analysis with immune gene platform was performed using mammary tumors obtained from Study 1 from mice fed high-MACi diet and treated with either IgG, anti-PD1, TAM, or TAM+anti-PD1. A total of 123 genes were differentially expressed in mice fed high-MACi diet and treated with TAM+anti-PD1 versus those treated with IgG, anti-PD1 or TAM monotherapy (**Supplementary Table 3**). In the KEGG analysis of these genes, several pathways were identified that were significantly different in TAM+anti-PD1 group, compared with the three other treatment groups. Of these, excluding any disease specific pathways, we focused on 12 relevant pathways in which more than 10% of pathway genes were differentially expressed. The highest percentile of differentially expressed genes (17.4%) belonged to the Th17 cell differentiation pathway. Others were Toll-like receptor signaling, NFκB signaling, PDL1 expression and PD1 checkpoint in cancer, AGE-RAGE signaling, JAK-STAT signaling, Th1 and Th2 differentiation, IL-17 signaling, HIF-1 signaling, TNF signaling, cytokine-cytokine receptor interaction and C-type lectin receptor signaling pathways, in the order of highest to lowest percentile of genes involved per pathway (**Table 2**).

**Table 2.** Signaling pathways identified in tumors of high-MACi mice treated with TAM+anti-PD1, compared IgG, TAM or anti-PD1.

| Pathway | % altered genes, of total genes in the pathway |
| --- | --- |
| TH17 cell differentiation | 17.4% |
| Toll-like receptor signaling | 16.5% |
| NFkB signaling | 16.2% |
| PDL1 expression and PD1 checkpoint in cancer | 15.6% |
| AGE-RAGE signaling in diabetic conditions | 13.9% |
| JAK-STAT signaling | 13.1% |
| Th1 and Th2 differentiation | 12.9% |
| IL-17 signaling | 11.6% |
| HIF-1 signaling | 11.0% |
| TNF signaling | 10.9% |
| Cytokine-cytokine receptor interaction | 10.7% |
| C-type lectin receptor signaling | 10.6% |

Significant different pathways were identified by KEGG analysis of NanoString data from mammary tumors of high-MACi fed mice treated with TAM+anti-PD1, compared with tumors in mice treated with IgG, TAM or anti-PD1. Only non-cancer pathways in which at least 10% of all genes in the pathway were differentially expressed in the analysis are included.

### Confirming gene expression changes in mammary tumors

To confirm differential gene expression within the Th17 differentiation pathway in E0771 tumors from mice fed high-MACi diet, we focused on *Batf*, *Jun*, *Rela*, *Rorα*, *Rorc*, and *Smad4*. Of these, in the TAM+anti-PD1 combination treatment group, *Rorα* and *Rorc* were upregulated compared with all other treatment groups (for p-values, see **Fig. 6A, B**), and *Batf* (p=0.030) **(Fig. 6C**) and *Rela* (p=0.037) (**Fig. 6D)** were suppressed compared with TAM. These findings are consistent with data obtained by NanoString. Changes in *Jun* or *Smad4* could not be confirmed **(Fig. 6E,F**).

**Figure 6.**
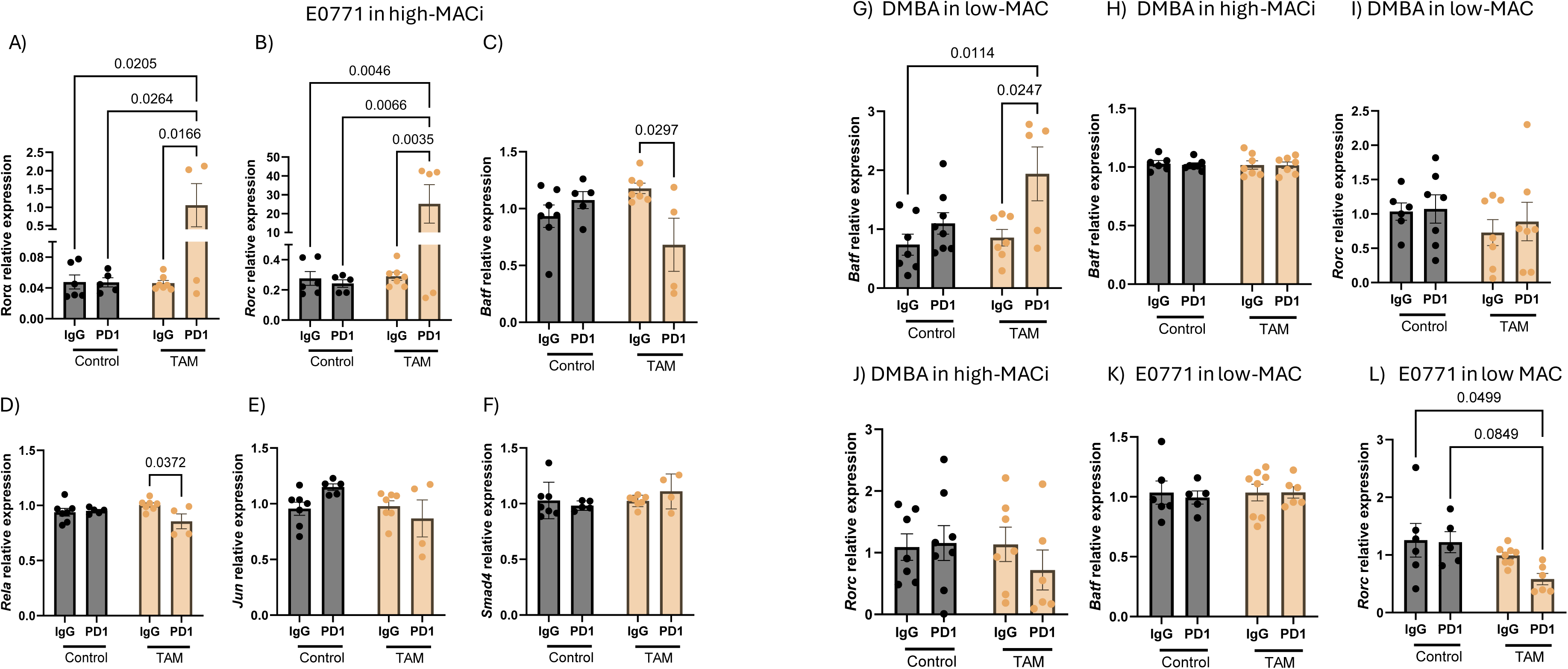
Confirmation of expression of genes in the Th17 differentiation pathway. Expression of (A) *Rorα*, (B) *Rorc*, (C) *Batf*, (D) *Rela*, (E) *Jun*, and (F) *Smad4* in mammary tumors of mice fed high-MAC isoflavone (MACi) diet and treated with IgG, anti-PD1, tamoxifen (TAM) or TAM+ anti-PD1. Expression of (G,H) *Batf* and (I,J) *Rorc* in 7,12-dimethylenz[a]anthracene (DMBA)-initiated estrogen receptor α positive (ERα+) mammary tumors in low-MAC or high MACi diet, or (K) *Batf* and (L) *Rorc* in E0771 TNBCs in low-MAC fed mice. Mice were treated with IgG, anti-PD1, tamoxifen (TAM) or TAM+anti-PD1. There were 4-8 tumors in each group. Data are shown as mean ± SEM with statistical significances obtained by two-way ANOVA followed by Tukey’s multiple comparisons test.

We next investigated whether Th17 differentiation pathway genes *Batf* and *Rorc* were down-regulated also in ERα+ mammary tumors in mice fed low-MAC diet, since combination treatment with TAM+anti-PD1 converted ICB unresponsive ERα+ tumors responsive in mice fed this diet. *Batf* was significantly upregulated in low-MAC diet fed mice treated with TAM+anti-PD1 compared with untreated control mice (p=0.01) or TAM (p=0.02) treated mice (**Fig. 6G**), but there were no differences among groups of mice fed high MACi (**Fig. 6H**). The expression of *Rorc* among mice treated with IgG, anti-PD1, TAM or their combination in ERα+ model in mice fed low-MAC or high-MACi diet was not affected either (**Fig. 6I,J**). In the TNBC model in mice fed low-MAC diet, *Batf* expression was not altered among the four treatment groups (**Fig. 6K**), but combination treatment (TAM+anti-PD1) reduced *Rorc*, compared with IgG treated mice (p=0.05) (**Fig. 6L**). Thus, up-regulation of *Rorα* and *Rorc* and downregulation of *Batf* and *Rela* may be linked to reversal of ICB resistance in TNBC model, but not in ERα+ model. In the ERα+ mammary tumors, upregulation of *Batf* was associated with response to anti-PD1 therapy under low MAC diet and co-treatment with TAM. Taken together, our findings indicate that although

Th17 differentiation pathway might be related to improved anti-PD1 response in TAM treated mice, individual genes in this pathway driving responsiveness seem to be different in TNBC and ERα+ mammary tumor models.

## Discussion

The gut microbiome plays an important, yet not clearly defined role in determining effectiveness of ICB (33, 34). Diet, especially fiber, modifies the gut microbiome (35), but studies have not consistently shown that fiber intake promotes response to ICBs (5). We studied here whether the effectiveness of anti-PD1 therapy in TNBC and ERα+ mammary tumor models that have not been included into previous studies investigating connections between dietary fiber and ICB response, might be different depending on whether mice are fed low– or high-MAC diet. The diets in our study contained different amounts isoflavones. Some evidence indicates that fiber improves the response to ICB therapy (1), while isoflavones may either improve or impair the response (8). Our results indicated that both low– and high-MAC diet fed mice responded to anti-PD1 therapy, but if mice were fed high-MAC diet containing notable levels of isoflavones, they did not.

Analysis of fecal SCFA levels showed that high-MAC and high-MACi diets significantly increased fecal acetic, butyric, propionic and valeric acids compared with low-MAC diet, with the high-MACi diet resulting in the highest increases (3-10-fold higher SCFAs). Other studies have found an increase in fecal SCFAs in mice fed high MAC diet, compared with low MAC diet like the one used in our study (36, 37). These results suggest that elevated fecal SCFA levels alone do not determine responsiveness to cancer immunotherapies. At the gut microbial level, both high-MAC and high-MACi diets elevated alpha-diversity and the abundance of SCFA producing bacterial families *Lachnospiraceae* and *Oscillospiraceae*, compared with low-MAC diet. Low-MAC diet elevated the abundance of SCFA producing *Muribaculaceae*, and high-MAC diet, but not high-MACi diet, increased the abundance of *Muribaculum* genus of *Muribaculaceae* family. These results suggest that diets which resulted responsiveness to anti-PD1 therapy elevated either fecal *Muribaculaceae* or *Muribaculum*. Similar data were generated in a human study which showed that high fecal abundance of *Muribacum* correlated with increased effectiveness of cancer immunotherapy (38).

Since high-MACi diet fed mice, which had the highest fecal SCFA levels, did not respond to anti-PD1 monotherapy, we studied whether isoflavones might blunt the response to anti-PD1. Isoflavone genistein binds and activates ERα and ERβ (26), the latter more than the former (28), and breast cancer cells that express higher levels of ERβ than ERα are inhibited by genistein (26). Genistein did not directly stimulate the growth of murine E0771 cells *in vitro,* consistent with E0771 mammary tumor cells expressing ERβ but not ERα (39), as also confirmed here. In immune cells, activation of ERα induces immunosuppression, while activation of ERβ stimulates anti-tumor immunity (40). However, since T cells in the TME express higher levels of ERα than ERβ (41), genistein may have promoted immunosuppression, explaining the failure of E0771 tumors to respond to anti-PD1 *in vivo* when mice were fed high-MACi diet or low-MAC diet supplemented with genistein.

ERα antagonist TAM dramatically improved anti-PD1 responsiveness in mice fed high-MACi diet, possibly because TAM has been reported to activate CD8+ T cells (30) in the TME. In earlier studies, another ERα antagonist (fulvestrant) improved response to ICBs in estrogen insensitive and ICB resistant preclinical non-breast cancer models by inhibiting immunosuppressive MDSCs (9, 14). We did not assess the effects of genistein on immune cells, but only focused on CD8+ T cell activation and exhaustion in mice fed low– or high-MAC/MACi diets. In human studies, when breast cancer TME contains high levels of exhausted CD8+ T cells, patients are most responsive to ICB (42, 43). In our study, CD8+ T cell infiltration into E0771 TME was similar in the high-MAC and high-MACi diet fed mice, but non-significantly lower in low-MAC diet fed mice. However, the percentage of exhausted CD8+ T cells was highest among mice fed high-MAC diet, and treatment with anti-PD1 significantly reduced an exhaustion marker in these cells. Anti-PD1 treatment tended to have an opposite effect on high-MACi diet fed mice, consistent with high-MACi diet fed mice not responding to anti-PD1 monotherapy. Our results thus agree with exhausted CD8+ T cells being a marker of responsiveness to ICBs in TNBC. Among low-MAC diet fed mice which exhibited very low levels of exhausted CD8+ T cells but also responded to anti-PD1, anti-PD1 treatment increased total CD8+ T cell infiltration into the TME. Future studies need to address in more detail the effect of genistein, alone or in combination with TAM, on immune cells in the TME.

Like ERα+ breast cancers in women (44) and animal models (45), ERα+ DMBA mammary tumors in our study were resistant to ICB. However, DMBA tumors became responsive to anti-PD1 treatment if mice were fed isoflavone-free low-MAC diet and co-treated with TAM. Whether patients with advanced ERα+ breast cancer might be more likely to respond to anti-PD1 therapy if they consume diets low in estrogens should be studied. High estrogenic foods are those that elevate circulating estrogen levels or that contain estrogenic compounds, like isoflavones. Foods that elevate circulating estrogen levels have been linked to increased breast cancer risk (46, 47), but their possible association to responsiveness to ICBs has not been investigated.

To study interaction between TAM and anti-PD1 in gene signaling level, we performed NanoString analysis using the cancer immune profiling platform. The Th17 cell differentiation pathway was the top immune signaling pathway that differed significantly in mice fed a high-MACi diet and were treated with a combination of TAM+anti-PD1, compared with the other three treatment groups. The role of Th17 cells in carcinogenesis has been conflicting (48). The controversy might be caused by the identification of two subtypes of Th17 cells: pathogenic and non-pathogenic (49). In our study, *RorA* and *RorC* were up-regulated and *Batf* and *Rela* were down-regulated in mice fed high-MACi diet and treated with TAM+anti-PD1. Upregulated *RorA* which is activated down-stream of IL6 to induce differentiation of Th17 cells (50), can act as a tumor suppressor gene (51) and is anti-inflammatory by inhibiting NFkB (52). The other upregulated gene, plays a key role downstream of IL6 and TGFβ, and works synergistically with *Rorα* to induce lineage specification of uncommitted CD4+ T cells into Th17 cells (53). *RorC* also improves responsiveness to cancer immunotherapy, including to anti-PD1 treatment (54). *Rela* promotes cancer cell growth and is involved in the polarization of uncommitted CD4+ T cells towards pathogenic Th17 cells (55). *Batf* also participates in maintaining the differentiation and function of Th17 cells (56). Down-regulation of *Rela* and *Batf* in high MACi fed mice treated with TAM+anti-PD1 is thus consistent with inhibiting of E0771 tumor growth.

Expression of *RorC* was not affected in ERα+ mammary tumors by diet or anti-PD1, TAM or TAM+anti-PD1 treatments. Instead, *Batf* was upregulated in low-MAC diet fed mice treated with TAM+anti-PD1 and consequently exhibiting reduced tumor growth. These results suggests that in the ERα+ mammary tumor model, *Batf* might have functioned to improve responsiveness to ICB treatment, as it has been reported to do (57). In summary, the combination treatment might have activated non-pathogenic Th17 differentiation in mice fed high MACi diet.

Clinical trials in breast cancer patients are on-going to determine the potential role of endocrine therapies in improving responsiveness to ICBs (58, 59). Whether the diet which these patients consume influences as to who benefits from the combination treatment has not been investigated. Our study highlights the importance of diet in determining ICB responsiveness in preclinical animal models. The results rebuff the idea that high MAC diets which elevate fecal SCFA levels always generate a better response to anti-PD1. Rather, the results indicate that even more important might be whether diet contains estrogenic components, such as isoflavones. Our findings show that isoflavones suppress response to anti-PD1, regardless of fecal SCFA levels, and blocking ERα+ in TNBC, i.e., probably in tumor microenvironment with TAM reversed the suppression. It is therefore crucial that when researchers are designing preclinical models to study cancer therapeutic response, they must consider the constituents of the laboratory diet of their animals and then report which lab chow was used. This will allow for accurate interpretation of results and ensure that the design of clinical trials based on these studies is appropriate. Our data also further highlights that human studies assessing the effectiveness of various combination therapies which include ICBs may need to consider inter-individual variation in patients’ diet.

## Supplementary Information

Randomization

Supplementary Table 1. DOCX. Composition of low-MAC, high-MAC and high MACi diets fed to mice.

Supplementary Table 2. DOCX. Primers used in quantitative real-time PCR.

Supplementary Table 3. DOCX. The list of 123 genes that differentially expressed in mice fed high MAC isoflavone diet and treated with TAM+anti-PD1 versus those treated with IgG, anti-PD1 or TAM.

## Declarations

### Ethics approval and consent to participate

The work described in the manuscript involves using preclinical models and in vitro experiments. The animal studies were reviewed and approved by Georgetown University and University of Minnesota Institutional Animal Care and Use Committees. All experiments were performed in accordance with the approved protocol and followed other relevant guidelines and regulations.

### Consent for publication

All authors have consented for publishing the manuscript

### Availability of data and material

The raw microbiome sequencing data that supports the findings of this study are openly available in the NCBI Sequence Read Archive (SRA) under BioProject accession number SRP579135. Other datasets that support the conclusions of this study are available from the corresponding author (L.H.C.) upon reasonable request.

### Authors’ contributions

*Conception and design*: L. Hilakivi-Clarke, P. Foley, F. de Oliveira Andrade

*Development of methodology*: F. de Oliveira Andrade, K. Andrade, V. Verma, K. Bouker, P. Foley, L.Hilakivi-Clarke

*Acquisition of data*: K. Bouker, I. Cruz, F. de Oliveira Andrade, K. Andrade, M Ozgul-Onal, A. Gao, C. Staley,

*Analysis and interpretation of data*: L. Jin, F. de Oliveira Andrade, K. Andrade, C. Staley, L. Hilakivi-Clarke,

*Writing the manuscript*: L. Hilakivi-Clarke, F. de Oliveira Andrade, C. Staley, K. Bouker, P. Foley

*Administrative, technical, or material support*: I. Cruz, W. Helferich

*Study supervision*: L. Hilakivi-Clarke

## Acknowledgements

Authors wish to thank Carlos Benitez and Alan Zwart at Georgetown University Medical Center for technical help in performing animal studies, and Dr. Esther Molino for technical support in performing in vitro experiments.

## List of abbreviations

ASVs: Amplicon sequence variants
BCAAs: branched-chain amino acids
BCFAs: branched-chain fatty acids
BSA: Bovine serum albumin
DMBA: 7,12-dimethylbenz[a]anthracene
DMSO: Dimethyl sulfoxide
E2: 17β-estradiol
ER: estrogen receptor
FBS: Fetal bovine serum
FMO: fluorescence minus one
GC-MS: gas chromatography-coupled mass spectrometry
GEN: genistein
GSEA: Gene Set Enrichment Analysis
High-MACi: high-MAC isoflavone
ICB: immune checkpoint blockade
MAC: microbiota-accessible carbohydrates
MPA: medroxyprogesterone acetate
PCoA: ordination using principal coordinate analysis
PD1: programmed cell death protein 1
SCFA: short-chain fatty acid
TAM: Tamoxifen
TBST: Tris-buffered saline with Tween 20
TME: tumor microenvironment
TNBC: triple negative breast cancer
UMGC: University of Minnesota Genomics Center

## References

1. Spencer CN, McQuade JL, Gopalakrishnan V, McCulloch JA, Vetizou M, Cogdill AP, et al. Dietary fiber and probiotics influence the gut microbiome and melanoma immunotherapy response. Science. 2021;374(6575):1632–40.

2. So D, Whelan K, Rossi M, Morrison M, Holtmann G, Kelly JT, et al. Dietary fiber intervention on gut microbiota composition in healthy adults: a systematic review and meta-analysis. Am J Clin Nutr. 2018;107(6):965–83.

3. Nomura M, Nagatomo R, Doi K, Shimizu J, Baba K, Saito T, et al. Association of Short-Chain Fatty Acids in the Gut Microbiome With Clinical Response to Treatment With Nivolumab or Pembrolizumab in Patients With Solid Cancer Tumors. JAMA Netw Open. 2020;3(4):e202895.

4. Luu M, Riester Z, Baldrich A, Reichardt N, Yuille S, Busetti A, et al. Microbial short-chain fatty acids modulate CD8+ T cell responses and improve adoptive immunotherapy for cancer. Nature Communications. 2021;12(1):4077.

5. Roichman A, Reyes-Castellanos G, Chen Z, Chen Z, Mitchell SJ, MacArthur MR, et al. Dietary Fiber Lacks a Consistent Effect on Immune Checkpoint Blockade Efficacy Across Diverse Murine Tumor Models. Cancer Res. 2025;85(17):3335–47.

6. Sonnenburg ED, Sonnenburg JL. Starving our microbial self: the deleterious consequences of a diet deficient in microbiota-accessible carbohydrates. Cell Metab. 2014;20(5):779–86.

7. Koh A, De VF, Kovatcheva-Datchary P, Backhed F. From Dietary Fiber to Host Physiology: Short-Chain Fatty Acids as Key Bacterial Metabolites. Cell. 2016;165(6):1332–45.

8. Focaccetti C, Izzi V, Benvenuto M, Fazi S, Ciuffa S, Giganti MG, et al. Polyphenols as Immunomodulatory Compounds in the Tumor Microenvironment: Friends or Foes? Int J Mol Sci. 2019;20(7).

9. Svoronos N, Perales-Puchalt A, Allegrezza MJ, Rutkowski MR, Payne KK, Tesone AJ, et al. Tumor Cell-Independent Estrogen Signaling Drives Disease Progression through Mobilization of Myeloid-Derived Suppressor Cells. Cancer Discov. 2017;7(1):72–85.

10. Chakraborty B, Byemerwa J, Shepherd J, Haines CN, Baldi R, Gong W, et al. Inhibition of estrogen signaling in myeloid cells increases tumor immunity in melanoma. The Journal of clinical investigation. 2021;131(23):e151347.

11. Milette S, Hashimoto M, Perrino S, Qi S, Chen M, Ham B, et al. Sexual dimorphism and the role of estrogen in the immune microenvironment of liver metastases. Nat Commun. 2019;10(1):5745.

12. Tai P, Wang J, Jin H, Song X, Yan J, Kang Y, et al. Induction of regulatory T cells by physiological level estrogen. J Cell Physiol. 2008;214(2):456–64.

13. Ye Y, Jing Y, Li L, Mills GB, Diao L, Liu H, et al. Sex-associated molecular differences for cancer immunotherapy. Nat Commun. 2020;11(1):1779.

14. Marquez-Garban DC, Deng G, Comin-Anduix B, Garcia AJ, Xing Y, Chen HW, et al. Antiestrogens in combination with immune checkpoint inhibitors in breast cancer immunotherapy. J Steroid Biochem Mol Biol. 2019;193:105415.

15. Gohl DM, Vangay P, Garbe J, MacLean A, Hauge A, Becker A, et al. Systematic improvement of amplicon marker gene methods for increased accuracy in microbiome studies. Nat Biotechnol. 2016;34(9):942–9.

16. Schloss PD, Westcott SL, Ryabin T, Hall JR, Hartmann M, Hollister EB, et al. Introducing mothur: open-source, platform-independent, community-supported software for describing and comparing microbial communities. Appl Environ Microbiol. 2009;75(23):7537–41.

17. Staley C, Kaiser T, Vaughn BP, Graiziger CT, Hamilton MJ, Rehman TU, et al. Predicting recurrence of Clostridium difficile infection following encapsulated fecal microbiota transplantation. Microbiome. 2018;6(1):166.

18. Garcia-Villalba R, Gimenez-Bastida JA, Garcia-Conesa MT, Tomas-Barberan FA, Carlos Espin J, Larrosa M. Alternative method for gas chromatography-mass spectrometry analysis of short-chain fatty acids in faecal samples. J Sep Sci. 2012;35(15):1906–13.

19. Adams KJ, Pratt B, Bose N, Dubois LG, St John-Williams L, Perrott KM, et al. Skyline for Small Molecules: A Unifying Software Package for Quantitative Metabolomics. J Proteome Res. 2020;19(4):1447–58.

20. Loveland BE, McKenzie IF. Cells mediating graft rejection in the mouse. II. The Ly phenotypes of cells producing tumor allograft rejection. Transplantation. 1982;33(2):174–80.

21. Sagiv-Barfi I, Czerwinski DK, Levy S, Alam IS, Mayer AT, Gambhir SS, et al. Eradication of spontaneous malignancy by local immunotherapy. Sci Transl Med. 2018;10(426).

22. Ozdemir BC, Sflomos G, Brisken C. The challenges of modeling hormone receptor-positive breast cancer in mice. Endocr Relat Cancer. 2018;25(5):R319–R30.

23. Andrade FO, Jin L, Clarke R, Wood I, Dutton M, Anjorin C, et al. Social Isolation Activates Dormant Mammary Tumors, and Modifies Inflammatory and Mitochondrial Metabolic Pathways in the Rat Mammary Gland. Cells. 2023;12(6).

24. Clarke KR. Non-parametric multivariate analyses of changes in community structure. Australian Journal of Ecology. 1993;18:117–43.

25. Chen IX, Newcomer K, Pauken KE, Juneja VR, Naxerova K, Wu MW, et al. A bilateral tumor model identifies transcriptional programs associated with patient response to immune checkpoint blockade. Proceedings of the National Academy of Sciences. 2020;117(38):23684–94.

26. Chang EC, Charn TH, Park SH, Helferich WG, Komm B, Katzenellenbogen JA, et al. Estrogen Receptors alpha and beta as determinants of gene expression: influence of ligand, dose, and chromatin binding. Mol Endocrinol. 2008;22(5):1032–43.

27. Barkhem T, Carlsson B, Nilsson Y, Enmark E, Gustafsson J-Å, Nilsson S. Differential Response of Estrogen Receptor α and Estrogen Receptor β to Partial Estrogen Agonists/Antagonists. Molecular Pharmacology. 1998;54(1):105–12.

28. Kuiper GG, Lemmen JG, Carlsson B, Corton JC, Safe SH, van der Saag PT, et al. Interaction of estrogenic chemicals and phytoestrogens with estrogen receptor beta. Endocrinology. 1998;139(10):4252–63.

29. Oner G, Broeckx G, Van Berckelaer C, Zwaenepoel K, Altintas S, Canturk Z, et al. The immune microenvironment characterisation and dynamics in hormone receptor-positive breast cancer before and after neoadjuvant endocrine therapy. Cancer Med. 2023;12(17):17901–13.

30. Hulskotter KJ, W.; Allnoch, L.; Hansmann, F.; Schmidtke, D.; Rohn, K.; Flugel, A.; Luhder, F.; Baumgartner, W.; Herder, V. Double-edged effects of tamoxifen-in-oil-gavage on an infectious murine model for multiple sclerosis. Brain Pathology. 2021;31(6):e12994.

31. Larsson AM, Roxå A, Leandersson K, Bergenfelz C. Impact of systemic therapy on circulating leukocyte populations in patients with metastatic breast cancer. Sci Rep. 2019;9(1):13451.

32. Huang B, Warner M, Gustafsson J. Estrogen receptors in breast carcinogenesis and endocrine therapy. Mol Cell Endocrinol. 2015;418 Pt 3:240–4.

33. Sivan A, Corrales L, Hubert N, Williams JB, Aquino-Michaels K, Earley ZM, et al. Commensal Bifidobacterium promotes antitumor immunity and facilitates anti-PD-L1 efficacy. Science. 2015;350(6264):1084–9.

34. McCulloch JA, Davar D, Rodrigues RR, Badger JH, Fang JR, Cole AM, et al. Intestinal microbiota signatures of clinical response and immune-related adverse events in melanoma patients treated with anti-PD-1. Nature Medicine. 2022;28(3):545–56.

35. Wastyk HC, Fragiadakis GK, Perelman D, Dahan D, Merrill BD, Yu FB, et al. Gut-microbiota-targeted diets modulate human immune status. Cell. 2021;184(16):4137–53 e14.

36. Tuck CJ, De Palma G, Takami K, Brant B, Caminero A, Reed DE, et al. Nutritional profile of rodent diets impacts experimental reproducibility in microbiome preclinical research. Sci Rep. 2020;10(1):17784.

37. Schipper L, Tims S, Timmer E, Lohr J, Rakhshandehroo M, Harvey L. Grain versus AIN: Common rodent diets differentially affect health outcomes in adult C57BL/6j mice. PLoS One. 2024;19(3):e0293487.

38. Zhang M, Wei Z, Wei B, Lai C, Zong G, Tao E, et al. Microbiota-derived urocanic acid triggered by tyrosine kinase inhibitors potentiates cancer immunotherapy efficacy. Cell Host & Microbe. 2025;33(6):915–31.e9.

39. Le Naour A, Rossary A, Vasson MP. EO771, is it a well-characterized cell line for mouse mammary cancer model? Limit and uncertainty. Cancer Med. 2020;9(21):8074–85.

40. Yuan B, Clark CA, Wu B, Yang J, Drerup JM, Li T, et al. Estrogen receptor beta signaling in CD8(+) T cells boosts T cell receptor activation and antitumor immunity through a phosphotyrosine switch. J Immunother Cancer. 2021;9(1).

41. Zhu B, Tse LA, Wang D, Koka H, Zhang T, Abubakar M, et al. Immune gene expression profiling reveals heterogeneity in luminal breast tumors. Breast Cancer Res. 2019;21(1):147.

42. Terranova-Barberio M, Pawlowska N, Dhawan M, Moasser M, Chien AJ, Melisko ME, et al. Exhausted T cell signature predicts immunotherapy response in ER-positive breast cancer. Nature Communications. 2020;11(1):3584.

43. Tietscher S, Wagner J, Anzeneder T, Langwieder C, Rees M, Sobottka B, et al. A comprehensive single-cell map of T cell exhaustion-associated immune environments in human breast cancer. Nature Communications. 2023;14(1):98.

44. Rugo HS, Delord JP, Im SA, Ott PA, Piha-Paul SA, Bedard PL, et al. Safety and Antitumor Activity of Pembrolizumab in Patients with Estrogen Receptor-Positive/Human Epidermal Growth Factor Receptor 2-Negative Advanced Breast Cancer. Clin Cancer Res. 2018;24(12):2804–11.

45. Perez-Lanzon M, Carbonnier V, Cordier P, De Palma FDE, Petrazzuolo A, Klein C, et al. New hormone receptor-positive breast cancer mouse cell line mimicking the immune microenvironment of anti-PD-1 resistant mammary carcinoma. J Immunother Cancer. 2023;11(6).

46. Harris HR, Bergkvist L, Wolk A. An estrogen-associated dietary pattern and breast cancer risk in the Swedish Mammography Cohort. Int J Cancer. 2015;137(9):2149–54.

47. Guinter MA, McLain AC, Merchant AT, Sandler DP, Steck SE. A dietary pattern based on estrogen metabolism is associated with breast cancer risk in a prospective cohort of postmenopausal women. Int J Cancer. 2018;143(3):580–90.

48. Zhao Y, Liu Z, Qin L, Wang T, Bai O. Insights into the mechanisms of Th17 differentiation and the Yin-Yang of Th17 cells in human diseases. Molecular Immunology. 2021;134:109–17.

49. Stockinger B, Omenetti S. The dichotomous nature of T helper 17 cells. Nature Reviews Immunology. 2017;17(9):535–44.

50. Kalim UU, Biradar R, Junttila S, Khan MM, Tripathi S, Khan MH, et al. A proximal enhancer regulates RORA expression during early human Th17 cell differentiation. Clinical Immunology. 2024;264:110261.

51. Xiong G, Wang C, Evers BM, Zhou BP, Xu R. RORα suppresses breast tumor invasion by inducing SEMA3F expression. Cancer Res. 2012;72(7):1728–39.

52. Oh SK, Kim D, Kim K, Boo K, Yu YS, Kim IS, et al. RORα is crucial for attenuated inflammatory response to maintain intestinal homeostasis. Proc Natl Acad Sci U S A. 2019;116(42):21140–9.

53. He S, Yu J, Sun W, Sun Y, Tang M, Meng B, et al. A comprehensive pancancer analysis reveals the potential value of RAR-related orphan receptor C (RORC) for cancer immunotherapy. Front Genet. 2022;13:969476.

54. Xia L, Tian E, Yu M, Liu C, Shen L, Huang Y, et al. RORγt agonist enhances anti-PD-1 therapy by promoting monocyte-derived dendritic cells through CXCL10 in cancers. Journal of Experimental & Clinical Cancer Research. 2022;41(1):155.

55. Lalle G, Lautraite R, Bouherrou K, Plaschka M, Pignata A, Voisin A, et al. NF-κB subunits RelA and c-Rel selectively control CD4+ T cell function in multiple sclerosis and cancer. J Exp Med. 2024;221(6).

56. Schraml BU, Hildner K, Ise W, Lee WL, Smith WA, Solomon B, et al. The AP-1 transcription factor Batf controls T(H)17 differentiation. Nature. 2009;460(7253):405–9.

57. Seo H, González-Avalos E, Zhang W, Ramchandani P, Yang C, Lio CJ, et al. BATF and IRF4 cooperate to counter exhaustion in tumor-infiltrating CAR T cells. Nat Immunol. 2021;22(8):983–95.

58. Sonnenblick A, Im SA, Lee KS, Tan A, Telli M, Strulov Shachar S, et al. 267P Phase Ib/II open-label, randomized evaluation of second– or third-line (2L/3L) atezolizumab (atezo) + entinostat (entino) in MORPHEUS-HR+ breast cancer (M-HR+BC). Annals of Oncology. 2021;32:S479.

59. Alaluf E, Shalamov MM, Sonnenblick A. Update on current and new potential immunotherapies in breast cancer, from bench to bedside. Front Immunol. 2024;15:1287824.

